# A synthetic bacterial information transfer system functions in the mammalian gut

**DOI:** 10.1101/308734

**Authors:** Suhyun Kim, S. Jordan Kerns, Marika Ziesack, Lynn Bry, Georg K. Gerber, Jeffrey C. Way, Pamela A. Silver

**Affiliations:** Dept of Systems Biology, Harvard Medical School, Boston, MA, 02115, USA; Dept of Molecular and Cellular Biology, Harvard University, Cambridge, MA, 02138, USA; Wyss Institute of Biologically Inspired Engineering, Harvard University, Boston, MA, 02115,USA; Dept of Pathology, Brigham and Women’s Hospital, Boston, MA, 02115, USA

**Keywords:** control systems, interspecies communication, gut microbiome, synthetic biology, quorum sensing, *luxI/luxR*

## Abstract

The gut microbiome is intricately involved with establishing and maintaining the health of the host. Engineering of gut microbes aims to add new functions and expand the scope of control over the gut microbiome. To create systems that can perform increasingly complex tasks in the gut with multiple engineered strains it is necessary to program communication among these bacteria in the gut. Towards this goal, we engineered an information transfer system for inter-cellular communication, using native gut *Escherichia coli* and attenuated *Salmonella enterica* serovar Typhimurium. Specifically, we have taken two genetic circuits-one for signaling from the quorum sensing system and the other for memory from the bacteriophage genetic switch–and integrated them into a robust system that can report on successful communication in the mammalian gut. Our system provides a basis for the construction of a programmable gut consortia as well as a basis for further understanding of bacterial interactions in an otherwise hard-to-study environment.

## Introduction

The gut microbiome has crucial roles in establishing and maintaining the health of the host, including the development of the immune system, nourishment of the gut epithelial cells, production of metabolites, neuromodulation, weight changes, and protection against pathogens (Ayres et al., 2012; Dodd et al., 2017; Everard et al., 2013; Kim et al., 2017; Ridaura et al., 2013; Sampson et al., 2016; Yano et al., 2015). These roles depend on the composition of the microbiome as well as the interactions among the microbes and with the host. A vast amount of information exists regarding the composition of the microbial populations within the mammalian gut (Clemente, et al., 2012 for review; Peterson, et al., 2009). Recent studies have also begun to map locations and interactions within the microbiome through imaging analysis (Earle et al., 2015; Mark Welch et al., 2017; Whitaker et al., 2017). However, our understanding of the complex dynamics of bacterial interactions in the gut is still at an early stage.

The gut microbiome can be reshaped through various measures, such as dietary changes, antibiotic treatments, and fecal transplants (Smillie et al., 2018; Turnbaugh et al., 2009; Zou et al., 2018). These approaches result in wholesale changes with limited levels of control and definition. Engineering of gut microbes poses an opportunity to overcome these limitations and expand the scope of control over the gut microbiome. This synthetic biology approach has shown progress in several ways thus far. First, engineered bacterial sensors can successfully detect their target molecules in the gut, such as orally administered anhydrotetracycline (ATC) and gut inflammatory markers, thiosulfate and tetrathionate (Daeffler, et al., 2017; Kotula et al., 2014; Riglar et al., 2017). Second, engineered bacteria with information recording systems using genetic switches can report on the state of the gut through fecal sample analysis (Kotula et al., 2014; Slauch et al., 2000; Mimee et al., 2016). Third, synthetic circuits can enable bacteria to produce beneficial molecules in the gut to improve the health of the host (Riglar and Silver, 2018 for review). Finally, suicide switches can enhance the biocontainment strategy of the engineered bacteria (Chan et al., 2015; Stirling et al., 2017). Overall, these approaches have shown advances in programing gut microbes with distinct functions for potential practical applications.

Although significant progress has been made in engineering the gut bacteria, little is known about programming interactions. Harnessing the ability of microbes to communicate with one another will lead to higher levels of control over the engineered gut consortia. Bacterial quorum sensing is a powerful natural system of intercellular signaling that may be repurposed for this task. Quorum sensing relies on the production and the detection of a signal molecule that can freely diffuse in and out of cells (Fuqua et al., 2001; Miller and Bassler, 2001 for review). The Gram-negative bacterial quorum sensing archetype consists of a *luxI*-homolog encoding the synthase of an acyl homoserine lactone (acyl-HSL) variety and a *luxR*-homolog encoding a transcription factor that activates the target genes when it is bound to a compatible acyl-HSL (Marketon et al., 2002; Piper et al., 1993; Rosemeyer et al., 1998). In nature, the *luxI/R*-type quorum sensing is used by many species of bacteria to distinguish their low and high-cell population density states and modulate group behaviors such as biofilm formation and virulence factor production (Engebrecht et al., 1983; Passador et al., 1993; Swift et al., 2001). Quorum sensing has been adopted to create novel functions that utilize signal production and detection as parts of cellular machineries. Examples include creating artificial cell patterning, logic gates, synchronization, pathogen inhibition, population density regulation, and programmed auto-lysis to deliver drugs to tumor cells (Basu et al., 2005; Brenner et al., 2007; Din et al., 2016; Hwang et al., 2016; Scott et al., 2017; Tabor et al., 2009; Tamsir et al., 2011).

For an acyl-HSL-based artificial communication system to function in a natural environment, ideally there should be little or no pre-existing acyl-HSL to interfere with the system. Interestingly, the presence of acyl-HSL has not been detected in the mammalian intestine, with one exception where mice were pre-infected with the pathogen *Yersinia enterocolitica* that produces acyl-HSL varieties (Dyszel et al., 2010; Smith et al., 2008). These observations suggest that the mouse gut may have little or no pre-existing acyl-HSL, but when acyl-HSL-producing bacteria are introduced, acyl-HSL can be detected. We therefore hypothesized that repurposing the *luxI/R*-type systems would allow us to create an effective communication system for the engineered bacteria in the gut.

To test this hypothesis, we created a synthetic information transfer system of *E. coli* and attenuated *S*. Typhimurium that integrates sensing, communication, and reporting. Overall, our system performs the following operations. When the tetracycline analog, ATC, is administered, the signaler strain produces *N*-(3-oxo-hexanoyl)-L-HSL (3OC6HSL); this acyl-HSL acts as the inter-cellular signal to alert the responder strain that the signaler strain has received the primary signal. To “capture” the evidence of successful communication between the two strains in the gut, we programmed the responder to switch ON a transcriptional genetic switch that has been shown to be robust and genetically stable (Kotula et al., 2014; Riglar et al., 2017) upon detecting 3OC6HSL (Fig 1A, B). This genetic switch is modeled after the CI-Cro bi-stable switch of the bacteriophage lambda and once activated by the signal, retains the ON state even after the signal is removal (Kotula et al., 2014). The advantage of adopting such a genetic switch that retains “memory” is that it can report the evidence of successful information transfer even if it was transient or occurred early during the transit trough the gut.

**Fig 1.**
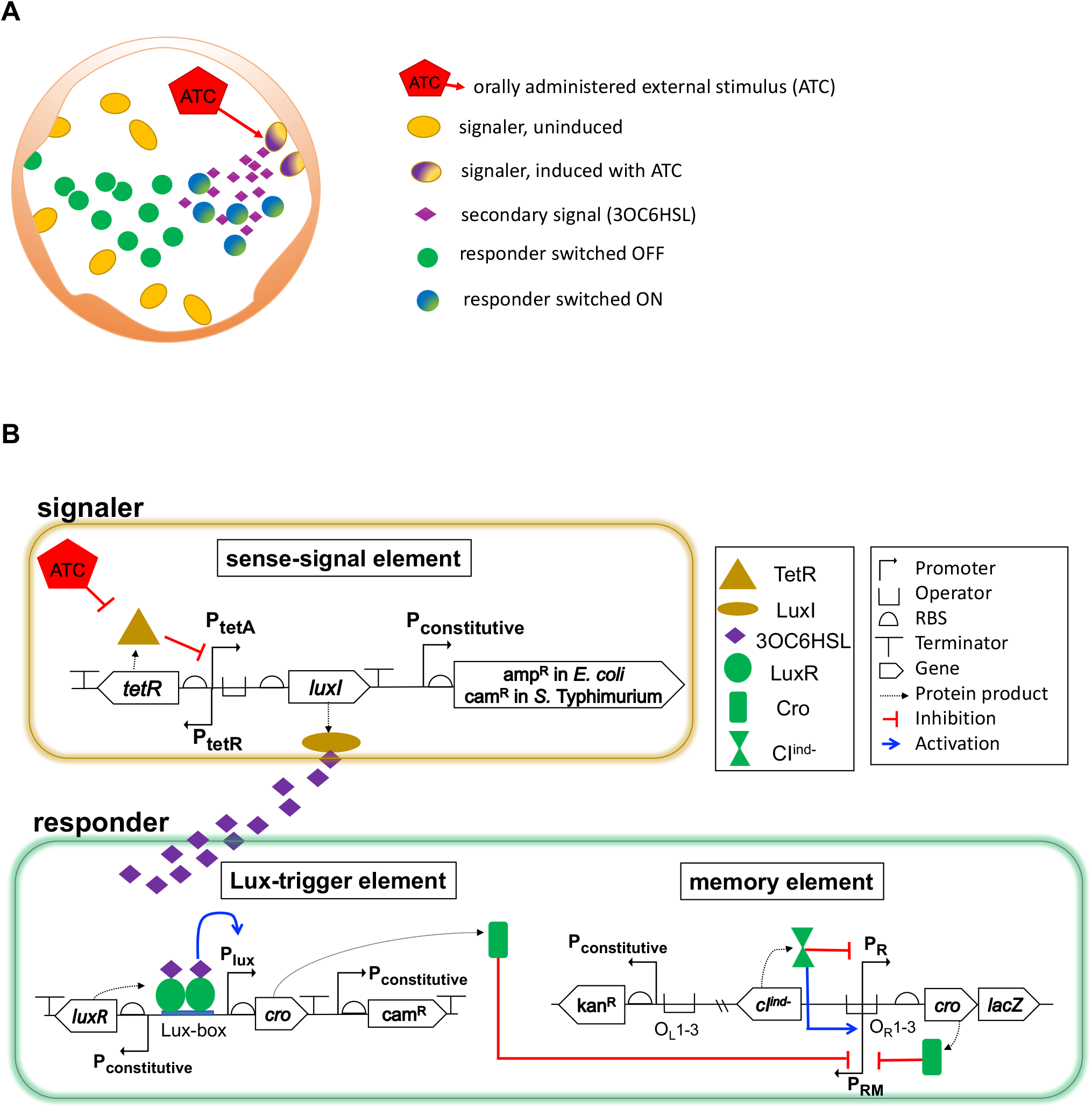
The information transfer system as a control element for the synthetic gut microbiome. (A) The abstraction of the information transfer system on the cross section of the gut. The signaler responds directly to the primary signal, ATC, and propagates the information to the responder using the secondary signal, 3OC6HSL. The responder switches ON in response to 3OC6HSL. (B) The diagram of the genome-integrated single copy circuits to make the signaler and the responder sense, relay, and record information as a pair. The signaler uses the sense-signal element to express *luxI* in the presence of ATC. The responder uses the Lux-trigger to express *cro* in the presence of 3OC6HSL. Once enough Cro is accumulated, the memory element switches ON to express *lacZ* as well as the second copy of *cro* to maintain the ON state.

## Results

### Engineering the information transfer system in *E. coli* and *S*. Typhimurium

We designed genetic elements to integrate ATC sensing, acyl-HSL-dependent communication, and memory into one coherent system. The signaler was engineered to produce 3OC6HSL in response to an external stimulus, ATC, using what we termed the sense-signal element. In the absence of ATC, the sense-signal element constitutively produces Tet repressors (TetR) that inhibit the P*tetA* promoter from transcribing *luxI*, which encodes the synthase of 3OC6HSL. ATC de-represses the inhibitory effect and allows the production of 3OC6HSL (Fig 1B). The responder was engineered to detect and record the reception of 3OC6HSL through the Lux-trigger element and the memory element. The Lux-trigger element leads to the expression of cro when the responder detects 3OC6HSL (Fig 1B). This element constitutively produces the *luxR* protein, which binds to 3OC6HSL and interacts with the Lux-box element of the P*luxI* promoter to activate the transcription of *cro* (Fig 1B). The *Cro* produced from the Lux-trigger element then acts on the previously described memory element (Kotula et al, 2014) to switch to a stable ON state. Once the memory element is switched ON, two genes, *lacZ* and the second copy of *cro*, are expressed through the memory element (Fig 1B). The expression of *lacZ* enables the use of blue-white colony screening to identify the ON state responder cells. The expression of the second copy of *cro* through the memory element forms a positive feedback loop to maintain the ON state even when the first copy of *cro* in the Lux-trigger element is no longer expressed.

The signaler and the responder were first constructed in the commensal murine gut *E. coli* strain, NGF, which has been shown to stably colonize the mouse gut (Kotula et al., 2014). The signaler was engineered by integrating a copy of the sense-signal element into the NGF genome. Using the NGF strain already containing the memory element (Kotula et al., 2014), we constructed the responder by further integrating the Lux-trigger element via Tn7 integration system.

Next, we constructed the *S*. Typhimurium version of the information transfer system. We first generated an attenuated strain background without the SPI-1 and the SPI-2 pathogenicity islands, which are involved in the Type-III secretion system for invasion into the host cells. The resulting strain did not have any outwardly visible adverse effect on mice even when it colonized the gut at high densities (>10^9^ CFU/g of feces, Supp Fig 2A). The *S*. Typhimurium signaler and the responder were built with the same design principles as those used in engineering the *E. coli* equivalents (Fig 1B). The *S*. Typhimurium signaler was created by integrating the *S*. Typhimurium codon-optimized sense-signal element into the *yafB-yafC* intergenic locus, where incorporating a large synthetic DNA does not lead to an obvious growth defect. The *S*. Typhimurium responder was created by integrating the *S*. Typhimurium codon-optimized Lux-trigger element and the memory element, which was adopted from the *E. coli* responder.

**Fig 2.**
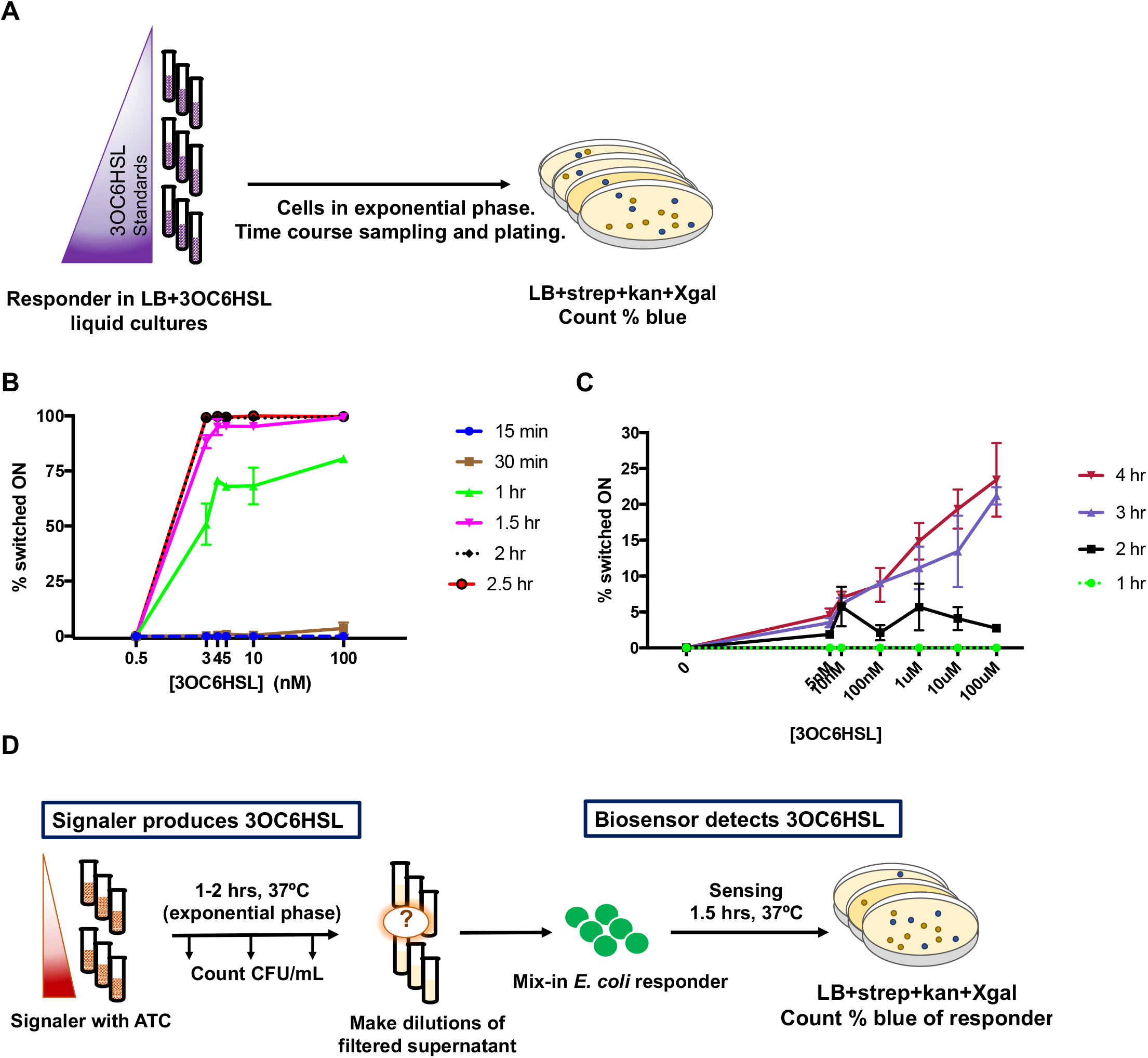
Characterization of the responder and the signaler as the biosensor and the producer of 3OC6HSL, respectively. (A) Experimental procedure to characterize the time course dosage response of the responder. The responder was sampled every 0.5h from t=0 to 2.5h (*E. coli*) or hourly from t=0 to 4h (*S*. Typhimurium) to count the % blue colonies. (B) *E. coli* responder sensing 3OC6HSL. Mean with SD; n=3 (C) *S*. Typhimurium responder sensing 3OC6HSL. Mean with SD; n=3. (D) Procedure to quantify the production rates of 3OC6HSL by the signalers. The signaler was induced with ATC (100ng/mL) for 1h (*E. coli*) or 2h (*S*. Typhimurium) in LB liquid media. CFU/mL values were obtained to obtain the signaler growth curve. To the serially diluted, cell-free signaler supernatants, the *E. coli* responder was mixed in for 1,5h for biosensing. Control responder cells were exposed to the 3OC6HSL standards to get the standard curve. % blue colony values were obtained by using Xgal plates.

### The *E. coli* and the *S*. Typhimurium responders switch ON in response to 3OC6HSL

We first characterized the responder as the biosensor of 3OC6HSL and subsequently used it to characterize the signaler as the producer of 3OC6HSL. The *E. coli* responder demonstrated a time-dependent dosage response to 3OC6HSL with high sensitivity and a good dynamic range. We exposed the *E. coli* responder to different concentrations of the 3OC6HSL standards for different lengths of time. The % responder switched ON was calculated by plating the samples on agar indicator plates containing Xgal and counting the % blue colonies (Fig 2A). The *E. coli* responder showed high sensitivity to the signal, with more than 50% of the cells switched ON with 3nM at t=1h. By t=2h, more than 98% of the responder cells switched ON, even with the lowest concentration tested (3nM) (Fig 2B). As expected, longer exposure time led to higher % switching ON.

The *S*. Typhimurium responder also demonstrated sensing of 3OC6HSL, with more cells switching ON at higher concentrations of 3OC6HSL and with longer exposure times (Fig 2A,2C). In comparison to the *E. coli* responder, however, it had lower sensitivity and a slower response rate; after exposure to the highest concentration of 3OC6HSL tested (100uM) for 4h, the *S*. Typhimurium responder switched ON to about 25% (Fig 2C).

### The *E. coli* and the *S*. Typhimurium signalers sense ATC and efficiently produce 3OC6HSL

Using the *E. coli* responder as a biosensor of 3OC6HSL, we quantified the production rates of 3OC6HSL by the *E. coli* and the *S*. Typhimurium signalers. The *E. coli* signaler demonstrated production of 3OC6HSL with high efficiency and capacity when induced with ATC. The signaler was induced with ATC (100ng/mL) for 1h. The signaler growth curve was obtained based on the colony forming units per mL (CFU/mL) of the signaler at t=0, 30, and 60min (details in Methods). At the endpoint, the filtered and cell-free signaler supernatants were serially diluted into fresh media, into which the responder was inoculated to detect 3OC6HSL (Fig 2D). The concentrations of 3OC6HSL in the supernatants were calculated by comparing the % switched ON of the responder when exposed to the diluted signaler supernatants versus the 3OC6HSL standards (Methods). Furthermore, using the estimated contributing signaler cell numbers over time, the production rate of 3OC6HSL was obtained as the number of molecules produced per signaler cell per min (Table 1 and Methods). The negative control using the uninduced signaler supernatant resulted in no switching ON of the responder as expected.

**Table 1.** The rates of production of 3OC6HSL by the *E. coli* and the *S*. typhimurium signalers. The concentration of 3OC6HSL in the supernatant was obtained using non-linear fitting of the standard curve and interpolating the values based on the % switched ON of the responder in the supernatant dilutions. The area under curve reported is for the signaler growth curve during the exposure to ATC. The production rate was obtained by dividing the total number of 3OC6HSL molecules in the supernatant by the area under growth curve. Mean ±SD of the multiple interpolated values from the signaler supernatant dilutions. (Details in Methods).

|  | [3OC6HSL] in supernatant (nM) | Area under curve | Calculated production rate (molecules made per cell per min) |
| --- | --- | --- | --- |
| *E. coli* | 27 ± 6.9 | 7.5 x 10<sup>8</sup> | 22000 ± 5500 |
| *S. Typhimurium* | 24 ± 2.6 | 3.5 x 10<sup>9</sup> | 4100 ± 440 |

The *S*. Typhimurium signaler also successfully produced 3OC6HSL when induced with ATC, with the efficiency and the capacity lower than those of the *E. coli* signaler. Using the same assay with minor modifications (Fig 2A), we found that compared to the *E. coli* signaler, the *S*. Typhimurium signaler produced approximately 6-fold less 3OC6HSL per cell per min in response to ATC (Table 1). Nevertheless, we expected that this production rate was high enough to lead to successful information transfer in the mouse gut, based on the observed density in the mouse gut for our attenuated strain of *S*. Typhimurium (>10^9^ CFU/g). For instance, the production rate of ~4100 molecules per signaler per min (Table 1) means that if the local density of the signaler in a closed environment is 5×10^8^ CFU/mL, after 1 min the concentration of 3OC6HSL would have already reached 3nM, which is approximately the EC50 value of the *E. coli* responder in our time course dosage experiment (Fig 2B, at exposure time 1h).

### Model describes the dynamic changes of the environment shaped by the signaler

To further our understanding of the system, we modeled how the signaler causes the dynamic changes in the concentration of 3OC6HSL in the immediate environment. The model presupposes three conditions: the environment is constantly mixing; the degradation of 3OC6HSL within the time scale of interest is negligible; and the net flux of 3OC6HSL in and out of the said environment is zero. In the case of constantly-mixing short-term *in vitro* cultures where these conditions are satisfied, the model is expected to provide a proper description of the system. In the case of more complex settings such as the gut, the model would serve as an approximate description of the immediate surrounding environment. The concentration of 3OC6HSL would increase in proportion to the density and the resident time (how long it has stayed in the given environment) of the signaler population (Fig 3A).

**Fig 3.**
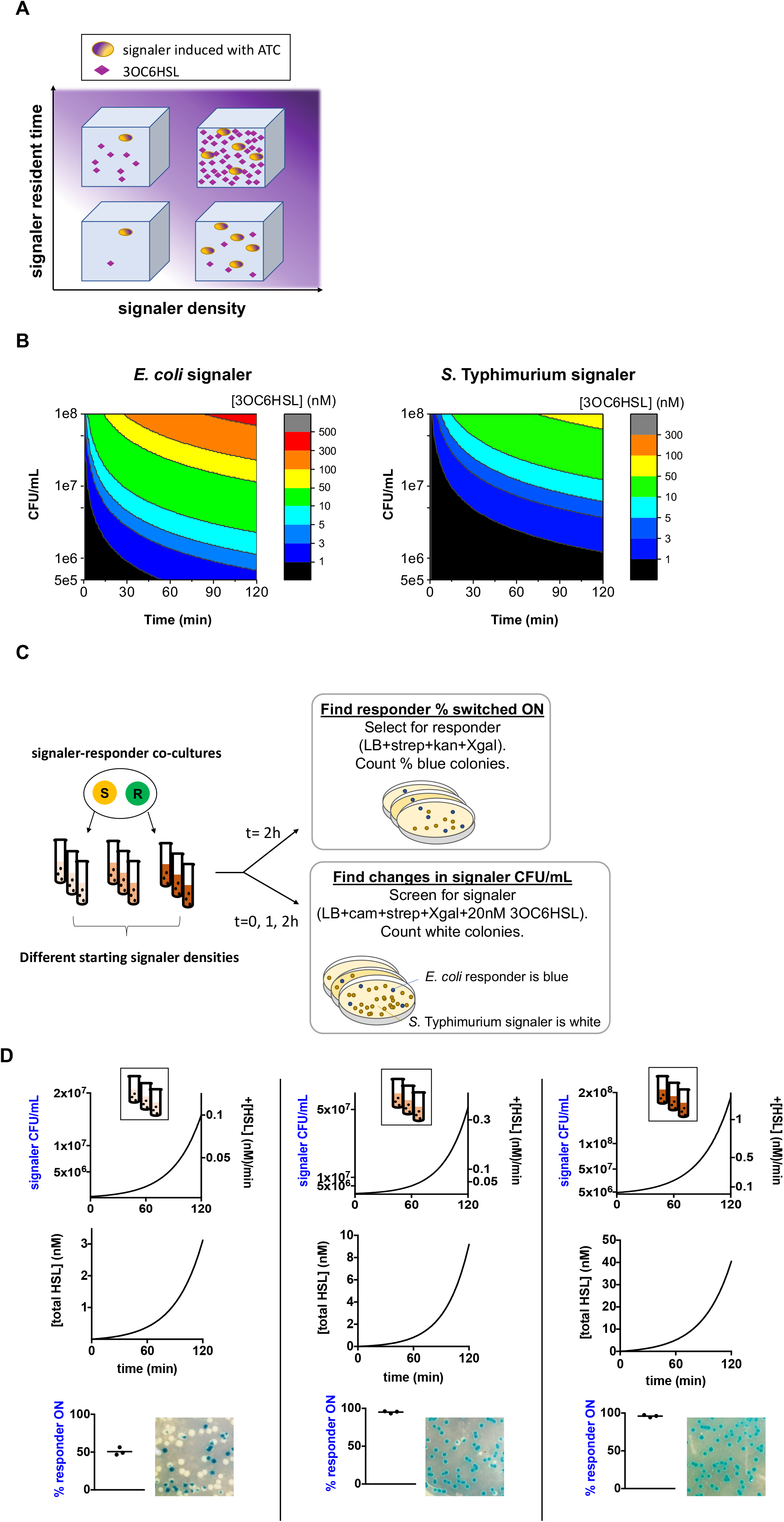
Model on the dynamic changes of the environment shaped by the signaler and its application to the *in vitro* co-cultures. (A) Abstraction on how the signal increases with higher signaler population density and the resident time in the microenvironment. (B) Predicted concentrations of the signal when the signaler population density is stable over time. The Y-axis denotes stable population densities and the heat map shows the predicted signal concentrations. (C) Procedure of the *S*. Typhimurium signaler-*E. coli* responder co-cultures. The cells were co-cultured with shaking in LB+ATC. Three sets had different starting signaler densities. The signaler CFU/mL values were obtained at T=0, 1, 2h to get the signaler growth curves. The responder % switched ON was determined at t=2h. (D) Each column corresponds to one set of triplicate co-cultures, with different starting signaler densities. From left to right: samples with increasing starting signaler densities. Blue axes were experimentally determined; black axes were obtained by applying the model to the experimentally determined signaler growth curves. Top row: experimentally determined signaler growth curves (left Y-axis) and the predicted signal production rates by the population (right Y-axis). Middle row: predicted total accumulated signal concentrations. Bottom row: experimentally determined % switched ON of the responder at t=2h. Bars indicate means. Pictures show the indicator plates for the responder blue-white screening.

The experimentally determined 3OC6HSL production rates of the *E. coli* and the *S*. Typhimurium signalers allowed quantitative predictions (Table 1, Fig 3B, 3D). If the signaler population density is at an equilibrium, the concentration of 3OC6HSL (nM) in the environment at any given time can be calculated as the product of the production rate ([HSL] (nM) produced per min per signaler), the population density (CFU/mL), and the resident time (min) of the signaler (Fig 3B). In cases in which the signaler population density evolves over time, the predicted concentration of 3OC6HSL (nM) at time t can be calculated by multiplying the production rate ([HSL] (nM) produced per min per signaler) to the antiderivative of the signaler density (CFU/mL) over t=0 to t (min).

Applying the model to the analysis of the *in vitro* co-cultures of the signaler and the responder demonstrates how it provides further understanding of the system. Three sets of the *S*. Typhimurium signaler-*E. coli* responder co-cultures with ATC were prepared, with different initial densities of the signaler inoculated (Fig 3C). The signaler growth curve was obtained based on the CFU/mL values of the signaler at t=0, 1, 2h. By applying the model, we could calculate the signal production rate by the population and the total accumulated concentration of the signal at any given time (Fig 3D). At the end of the co-culture (t=2h), the % switched ON of the responder was also determined by blue-white screening. One set with the lowest signaler density resulted in 50% of the responder switched ON and the other two sets with the higher signaler densities resulted in 95% of the responder switched ON (Fig 3D). As expected, the negative controls with the responder alone in ATC or the responder-signaler co-cultures without ATC remained OFF. The application of the model can thus complement the time course dosage response characterization of the responder (Fig 2A, 2B) by providing information on how the responder switches ON in conditions in which the concentration of the signal changes as the signaler population fluctuates (Fig 3D).

### The intra-species information transfer system in the mouse gut

The intra-species information transfer system deployed in the mouse gut demonstrated successful communication between the engineered *E. coli* signaler and the *E. coli* responder in a complex real-world environment. We delivered the two engineered strains into 12 conventional BALB/c mice by oral gavage (8 experimental, 4 control). Starting from two days after the gavage, the experimental group was given ATC (0.1mg/mL in drinking water) for two days. The control group was not given ATC at any point (Fig 4A). Each day, fecal samples were collected and analyzed on selective plates to count the CFU/g values of the signaler and the responder, as well as the % switched ON of the responder (Fig 4B, Methods). One day after the gavage and before ATC was given, the responder was OFF in all mice except one in the experimental group, which showed a low level (1%) of switching ON. After one day of dosing with ATC, five out of the eight mice in the experimental group showed clear signs of the responder switched ON, ranging from 13 to 74% in different mice. After two days on ATC, the same five mice still had the responder switched ON to various degrees (Fig 4C). There was an average 5-fold drop in the colonization of the responder between the days 0 and 3 (Supp Fig 1A). There was no clear correlation between the % switched ON and colonization level or the ratio between the two strains (Supp Fig 1B, 1C). The responder in the four mice in the negative control group without ATC remained OFF until the end of the study (Fig 4C). These data indicate successful information transfer between the signaler and the responder in the majority of mice.

**Fig 4.**
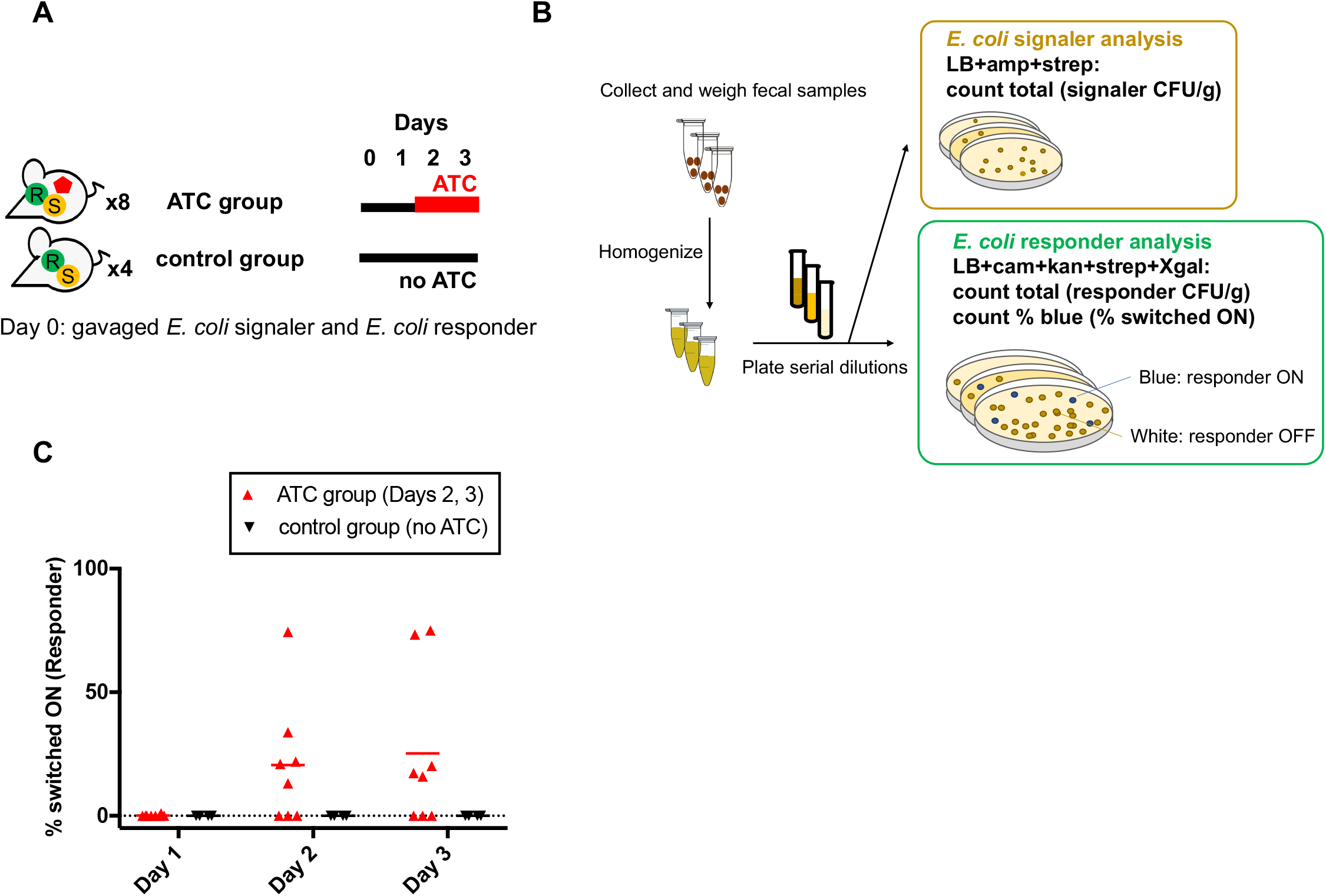
The *E. coli* to *E. coli* intra-species information transfer system in the gut. (A) Schematics of the animal study. (B) Downstream analysis of the daily fecal samples. Fecal samples from each mouse were weighed, homogenized, serially diluted, and plated onto selective plates for the **E. coli** signaler or the **E. coli** responder. (C) The switching ON of the responder in the ATC group and the control group. Bars indicate means.

### The inter-species information transfer system in the mouse gut using the *S*. Typhimurium signaler and the *E. coli* responder

The inter-species information transfer system consisting of the *S*. Typhimurium signaler and the *E. coli* responder also demonstrated successful communication in the mouse gut. We used the study design described above with minor modifications on the duration of the experiment and the selective plates used (Fig 5A, B). The result indicated that 7 out of 8 mice in the experimental group had the responder switched ON with ATC but not without ATC, with slight daily variations. The % switched ON ranged from 0.5% to 86% in different mice. (Fig 5C, Supp Fig 2C). All mice in the negative control group without ATC showed the responder in the OFF state throughout the experiment (Fig 5C). Unexpectedly, one mouse of the experimental group switched ON to around 50% one day before ATC was given (Fig 5C, Supp Fig 2C). To examine whether this switching ON was caused by mutations within the engineered parts, we sequenced the entire Lux-trigger and the memory elements of the switched ON responder cells in this mouse but found no mutation (see Discussion). From tallying the CFU/g each day, we did observe a significant drop in the *E. coli* responder densities throughout the study. The responder density dropped 30-fold on average by day 9, while the *S*. Typhimurium signaler maintained its density (Supp Fig 2A). Interestingly, the mouse with the unexpected high switching ON before ATC showed the most rapid and dramatic decrease in the responder population, with the density dropping more than 1000-fold by day 9 (Supp Fig 2A). There was no clear correlation between the % switched ON and the CFU/g or the ratio between the two species in other mice (Supp Fig 2B, 2C).

**Fig 5.**
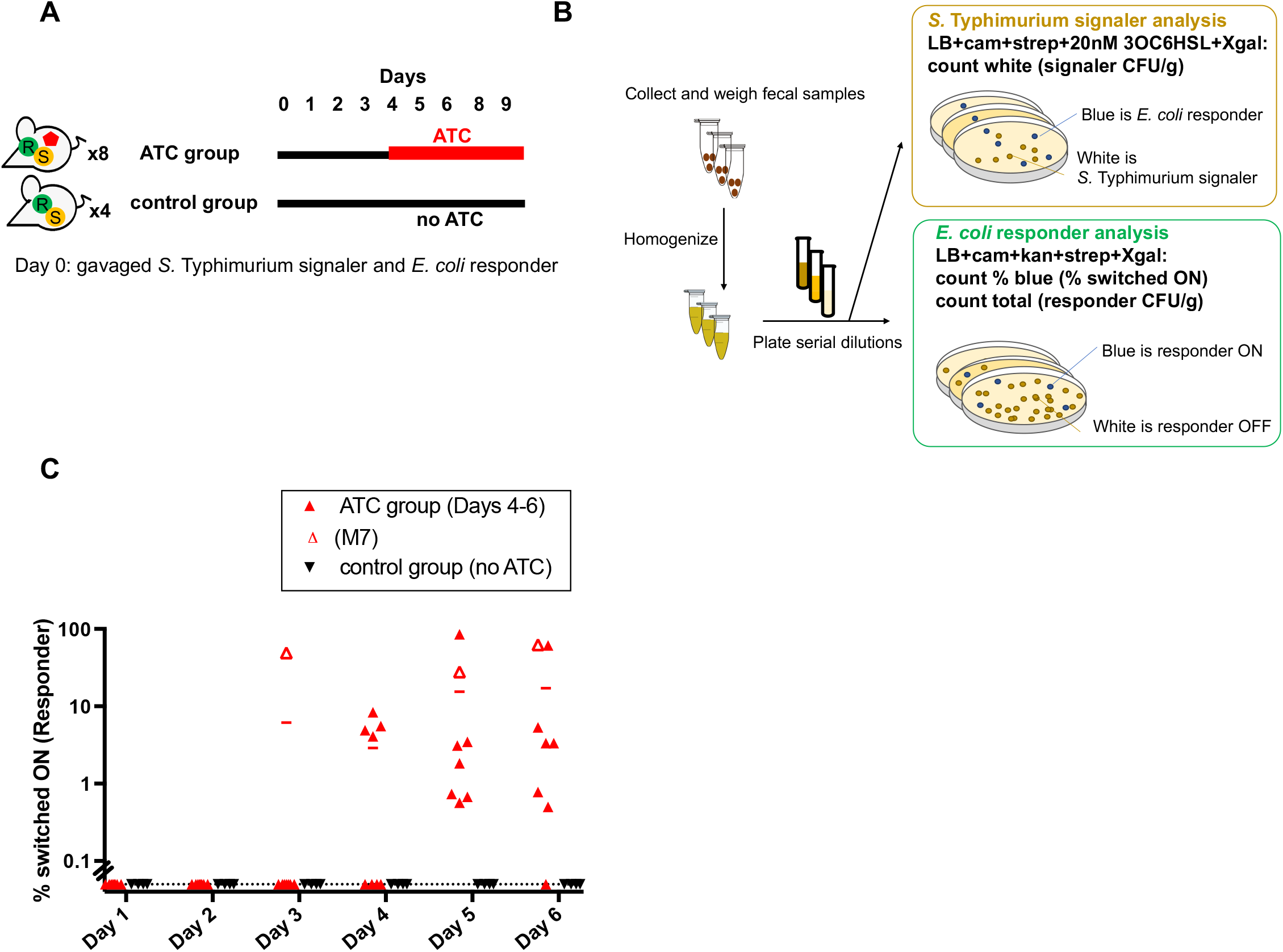
The *S*. Typhimurium to *E. coli* inter-species information transfer system in the gut. (A) Schematics of the animal study. The 8 mice in the experimental group received ATC after Day 3 in drinking water. The 4 mice in the control group were not given ATC. Fecal samples were collected for the downstream analysis. (B) Downstream analysis of the fecal samples. The addition of 20nM of 3OC6HSL in plates induces all **E. coli** responder cells to form blue colonies, while the *S*. Typhimurium signaler cells remain white. (C) The detected switching ON of the responder cells from the mouse fecal samples. M7 is the outlier that switched ON one day before ATC was given (see main text and Supp Fig 2). The % for M7 on Day 4 could not be counted due to an unexpected drop in the CFU/g. All data points below 0.1% ON were plotted together on the dotted line for simplicity. Bars indicate means.

## Discussion

Due to the close association of the gut micobiota with the mammalian host, it is attractive to consider engineering gut bacteria to monitor the health of the host, report on infections and diseases, and provide enhanced nutrition or treatments *in situ*. Programmable inter-microbial communication in the gut is an underexplored but key element to fully achieve these goals for the following reasons. To begin with, complex and extensive bacterial engineering can impose an increased metabolic burden on the cells that can lead to genetic instability and subsequent losses of the synthetic parts over generations (Borkowski et al., 2016; Sleight and Sauro, 2013; Wu et al., 2016). Division of labor among different microbes that can communicate with one another to coordinate their actions may provide one solution. Furthermore, programmable communication allows cells to form networks that can accomplish more complex tasks. The strengths of such networks of synthetic cells that coordinate their actions using secreted signals have been suggested in studies using *in vitro* systems (Ausländer et al., 2018; Tamsir, et al., 2011). These systems, consisting of networks of engineered bacteria or human cells, could accomplish computations and create cellular patterns that are more complex than the sum of the individual parts (Ausländer et al., 2018; Tamsir, et al., 2011). Similarly, we expect that the ability to program communication for the engineered gut consortia would allow more advanced tasks to be devised and performed *in vivo*.

Our study using the intra- and inter-species information transfer systems demonstrated successful communication among the gut bacteria using acyl-HSL as the inter-cellular signal. The overall design strategies included integrating all synthetic cassettes as genome-integrated single copies to increase stability and reduce the metabolic burden of the synthetic parts as well as incorporating a genetic switch for post-transit analysis through analysis of bacteria in fecal samples.

The *E. coli* responder demonstrated several strengths as an in vivo biosensor to record and report the evidence of communication between the signaler and the responder. First, the *E. coli* responder has high sensitivity to the signal of interest, 3OC6HSL; *in vitro* analysis showed that it has a low nM detection limit. Second, the responder switches ON after a physiologically achievable duration of exposure to the signal. When the responder cells in the exponential phase of growth were exposed to 3OC6HSL *in vitro*, the minimum required time of exposure for marked switching to occur was 1h, which is within the residency time of bacteria in the gut (Myhrvold et al., 2015). Finally, the responder serves a unique role in probing the gut, with a genetic switch-based memory element that can record even transient communication in the gut.

The *E. coli* and *S*. Typhimurium signalers demonstrated the production of 3OC6HSL with high efficiency and capacity when induced with ATC. Many *in vitro* synthetic systems have been built using acyl-HSLs as the inter-cellular signals among the engineered bacteria. However, these systems overexpress the *luxI* homologs from multi-copy plasmids and do not include quantification of the signal production. As such, it was unclear whether bacteria that do not naturally use quorum sensing can be engineered to produce physiologically effective levels of acyl-HSLs when produced by a single, genomically integrated copy of *luxI*. The rate of production by the signaler using the responder as the biosensor confirmed that it is possible. Furthermore, this rate of production was applied to build a model that predicts the dynamic changes in the concentration of the signal in each environment defined by the density and the resident time of the signaler.

Both the intra-and inter-species information transfer systems indicated that the signaler could successfully communicate to the responder using acyl-HSL in the gut, but also showed interesting complexities not observed *in vitro*. First, as with most animal experiments of this type, there was mouse-to-mouse variation in response to ATC. Although the majority of mice in the ATC group showed switching ON, a few mice remained OFF despite receiving ATC for a few days. We do not think this is due to reversion from the ON to the OFF state, since previous studies established the stability of the ON state (Kotula et al., 2014; Riglar et al., 2017). Alternatively, in the mice that showed no response, the signaler cells might not have been exposed to sufficient levels of ATC, either because these mice did not consume enough ATC-containing water, or because there are variations in the host metabolism of ATC. Second, among the mice that responded to ATC, the % switched ON of the responder cells varied. The degree of switching did not correlate with the CFU/g of the signaler or the ratio between the signaler and the responder. It is possible that the variation is caused by the differences in the background microbiota, the host metabolism, or the amount of ATC that successfully reached the gut in each mouse. Lastly, in our inter-species study, we observed an unexpected switching ON to ~50% without ATC in one mouse. Prior studies using the memory element over longer periods of up to 6 months have shown no mutation in the engineered bacteria that led to the untriggered ON state (Riglar et al., 2017). It is possible that the switching ON might have resulted from the bona fide detection of acyl-HSLs or agonists that were present.

The overall difference in the responder switching ON between the ATC-receiving group and the control group of mice demonstrated that communication can happen successfully using 3OC6HSL as the inter-cellular signal in the gut. The average % switched ON value was higher in the intra-species system than in the inter-species system (Fig 4C, 5C). This difference may be partly due to the production rate of 3OC6HSL being higher in the *E. coli* signaler than in the *S*. Typhimurium signaler (Table 1). It is also possible that the attenuated *S*. Typhimurium signaler colonizes at a further distance away from the *E. coli* responder than does the *E. coli* signaler, thus reducing the concentration of 3OC6HSL that the responder is exposed to.

In summary, we have integrated quorum sensing-based inter-cellular signaling and memory into the information transfer system that can function to report on cell interactions and communication in the mammalian gut. Our system provides a basis for further understanding of inter-bacterial interactions in an otherwise hard-to-study environment as well as a basis for the construction of programmable gut consortia.

## Methods

### Engineering the *E. coli* signaler and the responder

The *E. coli* signaler was engineered using the sense-signal element synthesized as a gBlock (IDT), with ~60 bp homology arms to target the intergenic region between *araB* and *araC*. The element has the sequence of Tn*10* from the 3’ end of tetR up until the start of *tetA. V. fischeri* MJ1 strain’s *luxI*, codon-optimized (IDT) for *E. coli*, was put in the place of tetA. 100ng of the gBlock was electroporated into the recombineering-competent strain of *E. coli*, TB10 (Johnson et al., 2004). Transformants were selected on LB+ampicillin 100ug/mL. In every recombineering step in this study, single colonies were picked and re-streaked twice before they were used in subsequent steps. The P1*vir* lysate from the transformed TB10 was used to transduce the NGF strain of *E. coli* to make the signaler. The *E. coli* NGF responder was made by introducing the Lux-trigger element into an existing NGF strain containing the memory element (Kotula et al., 2014). The Lux-trigger element was synthesized as a gBlock (IDT) with the following specifics. The *luxR* of the *V. fischeri* MJ1 was codon-optimized for *E. coli* and put downstream of the PlacF promoter and a synthetic RBS. The native *cro* gene from the bacteriophage lambda was put downstream of the *V. fischeri* MJ1’s P*luxI* promoter and a synthetic RBS (part number BBa_B0029, iGEM Registry of Standard Biological Parts). P*luxI* and P*lacI*^Q^ were separated by ~80 bp of random, buffering sequences. The gBlock was cloned into the pDR08 plasmid, a temperature-sensitive plasmid with arabinose-inducible genes for site-specific recombination using the Tn7 transposon elements. After growing the electroporated cells on plates with LB+10mM arabinose+12.5ug/mL chloramphenicol at 30°C for two days, single colonies were picked and passaged twice on LB+25ug/mL chloramphenicol at 42°C to cure the plasmids. The integration of the Lux-trigger element was confirmed via chloramphenicol resistance and the plasmid curing via ampicillin susceptibility.

### Engineering the *S*. Typhimurium signaler and the responder

To build an attenuated strain, we introduced a knockout of SPI-2 into the *S*. Typhimurium LT2 strain lacking the major part of SPI-1 (strain JS481, gift of Dr. James Slauch) (Ellermeier et al., 2005). The knockout of SPI-2 was first created in a recombineering-competent strain of LT2 using a gBlock to replace SPI-2 with a kanamycin-resistance cassette flanked by FRT sequences. The cassette was then P22-transduced into the strain JS481. A temperature-sensitive plasmid encoding FLPase was introduced to remove the kanamycin-resistance cassette. The plasmid was cured after confirming kanamycin sensitivity. The resulting, attenuated strain of LT2 was sequence-confirmed to be: Δ3009865-3044887 (of SPI-1) and Δ1501112-1462122 (of SPI-2), of the NCBI sequence ID AE006468.2). The sense-signal element and the Lux-trigger element were first integrated into the recombineering-competent LT2 strain (Lau et al., 2017) and was subsequently P22-transduced into the above attenuated strain background to make the signaler. The elements were synthesized as gBlocks, with 100 bp homology arms targeting the intergenic locus between *yafB* and *yafC*. The *S*. Typhimurium signaler resistant to streptomycin (>300ug/mL) was obtained by exposing cells to increasing concentrations of streptomycin and selecting for spontaneously arising resistant clones. To integrate the memory element to build the responder, we built the CRIM P21 integration plasmid and propagated it in the M+R-Salmonella strains DB7011 (gift of the David Botstein lab). The memory element was first integrated into the SA2009 strain from the Salmonella Genetic Stock Center and P22-transduced first into the LT2 strain and subsequently into the attenuated LT2 strain. The *S*. Typhimurium Lux-trigger element was incorporated into the recombineering-competent LT2 strain and P22-transduced into the attenuated strain harboring the memory element to generate the *S*. Typhimurium responder.

### Time course dosage responses of the *E. coli* and the *S*. Typhimurium responders

An overnight culture of the responder was diluted 1:4000 into LB+cam or LB+strep broth and grown for 1.5 hrs (E.coli responder) or 2.5 hrs (*S*. Typhimurium responder) to pass the lag phase and enter the exponential phase of growth. The culture was then divided into multiple tubes, to which 3OC6HSL (Sigma Aldrich, k3007; stock solutions were prepared using dimethyl sulfoxide and kept at −20°C) was added to various final concentrations. From each tube, 100uLx3 was distributed into three wells of a U-shaped, low-attachment 96-well plate for a triplicate reaction. Cells were grown at 37°C with shaking throughout the procedure. Between t=0-4h, a small volume from each well was taken out hourly and serially diluted in PBS to aim for ~10^3^−10^4^CFU/ml. 50-200uL were bead-plated onto LB+kan 50ug/mL+strep300ug/mL+Xgal 60ug/mL. The % blue colony values were counted.

### Quantification of the 3OC6HSL production by the *E. coli* and the *S*. Typhimurium signalers

The overnight culture of the signaler was subcultured in LB+strep broth for 1.5h (*E. coli*) or 2.5h (*S*. Typhimurium) to pass the lag phase and enter the exponential phase of growth. The signaler cells were then exposed to ATC (100ng/mL) for 1h (*E. coli*) or 2h (*S*. Typhimurium) to produce 3OC6HSL at 37°C with shaking. Small volumes were taken out and plated to count the CFU/g at t=0, 30min, 1h (*E. coli*) or t=0, 1h, 2h (*S*. Typhimurium) and obtain the growth curves. At the endpoint, cells were pelleted and the supernatants were sterile-filtered (0.2um). The supernatants were diluted 1:10, 20, 30, 40, 50 into LB+strep broth containing the pre-subcultured (~1.5h), post-lag phase *E. coli* responder in the exponential phase. The supernatant+responder mixtures were incubated at 37°C with shaking for 1.5h for the biosensor reaction. We countered possible day-to-day or batch-to-batch variations in the responder switching ON, by exposing the same batch of the responder cells to the 3OC6HSL standards in parallel, to serve as the reference dosage response of that day. At the end of the 1.5h exposure, the responder cells were diluted and plated onto LB+kan+strep+Xgal to count the % blue colonies. We used Graphpad Prism to interpolate the concentrations of 3OC6HSL of the diluted supernatants. First, we input the % blue colony values of the responder when exposed to different concentration of the 3OC6HSL standards for non-linear fitting. Then, we input the % blue colony values of the responder when exposed to the diluted signaler supernatants to obtain the interpolated concentrations of 3OC6HSL in these diluted supernatants. Multiplying these values by the dilution factors, we obtained the concentrations in the undiluted supernatants. These concentrations were used to calculate the rates of production of 3OC6HSL by the *E. coli* and the *S*. Typhimurium signalers using the following formula:

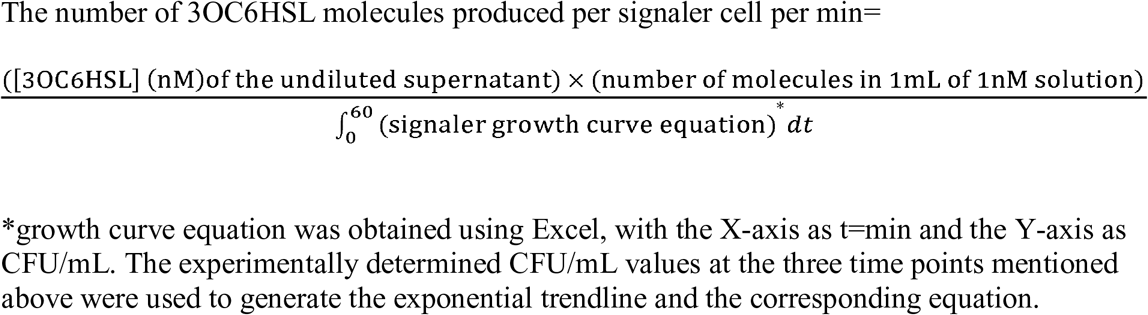

### *In vitro* co-cultures of the *S*. Typhimurium signaler and the *E. coli* responder

Overnight cultures (LB+strep 200ng/mL) of the attenuated *S*. Typhimurium signaler and the *E. coli* responder were subcultured (1:4000, LB+strep 100ng/mL) for 2.5h at 37°C. The signaler was then diluted 1/20, 1/10, and 1/2 (LB+strep 100ng/mL), to which the responder (1/500 of the subculture) and ATC (100ng/mL) was added and vortexed to mix. Two sets of negative controls were prepared: i) responder alone (1/500 dilution) with ATC (100ng/mL) and ii) responder (1/500 dilution) and the signaler (1/2 dilution) without ATC. This point was set as t=0 and each of the three co-culture sets and the negative controls was distributed into three wells of a U-shaped, low-attachment 96-well plate for triplicate reactions. The plate was incubated at 37°C on a plate shaker at ~1000rpm. At t=0, 1, 2h, small volumes of the samples were diluted in PBS and plated on LB+cam+strep+Xgal+20nM 3OC6HSL to screen for the *S*. Typhimurium signaler as white colonies. At t=2h, the samples were also plated onto LB+kan+strep+Xgal plates to count % blue colonies of the *E. coli* responder. The signaler growth curves were obtained using Excel by plotting exponential trendlines (X-axis as t=min; Y-axis as CFU/mL) and corresponding equations based on the experimentally determined CFU/mL values at t=0, 1, 2h.

### Intra-species information transfer in the mouse gut

The Harvard Medical School Animal Care and Use Committee approved all animal study protocols. Female BALB/c Elite mice of 6-week old (Charles River company) were acclimatized to the facility for 2 weeks before the start of experiments. Each cage housed four mice (M1-M4, M5-M8, and M9-M12) under the microisolator FST system in the BSL-1 facility. Mice were fed with regular high-fiber, low-fat diet (PicoLab Rodent Diet, 5053) and drinking water with strep (0.5mg/mL streptomycin sulfate (USP grade, VWR), 5% sucrose; 0.2uM sterile-filtered) starting from half a day before the gavage to the end of the experiment (Day 3). All 12 mice were given approximately 10^8^ CFU each of the *E. coli* signaler and the *E. coli* responder by oral gavage on Day 0. The experimental group (M1-M8) were provided with drinking water that contains ATC (0.1mg/mL of ATC (Sigma Aldrich), 0.5mg/ml strep, 5% sucrose, 0.2um filter-sterilized; water bottles were foil-wrapped due to ATC photosensitivity) during the treatment period (starting after the sample collection on Day 1). Special water was freshly made at least every 48h. Fecal samples were collected daily, weighed, suspended in PBS to 100mg/mL, and vortexed thoroughly to homogenize. The samples were centrifuged at low speed (100g, 15min) to pellet down large fibers and insoluble particles. The supernatant in the upper part of the tube was serially diluted (10^−3^−10^−6^-folds) in PBS and bead-plated (100uL on 10cm selective plates) for the *E. coli* signaler (LB+amp 50ug/mL+strep 300ug/mL) or the *E. coli* responder (LB+cam 25ug/mL+kan 50ug/mL+strep 300ug/mL+Xgal 60ug/mL). Samples collected before the oral gavage resulted in no growth on the selective plates, confirming the absence of the background flora resistant to the antibiotics used to select for the engineered cells.

### Inter-species information transfer in the mouse gut

The general information regarding the animal treatment and the sample collection is the same as described above, except the mice were acclimatized to the facility for 1 week and housed in the BSL-2 facility. Four mice were housed per cage (M1-4, M5-8, M9-12). Approximately 2×10^8^ CFU each of the *S*. Typhimurium signaler and the *E. coli* responder were delivered into all 12 BALB/c mice by oral gavage on Day 0. Drinking water with strep was given to all mice starting from 3 days before the gavage to the end of the experiment. The experimental group (M1-M8) were provided with drinking water with ATC during the treatment period (starting after the sample collection on Day 3). The signaler CFU/g was determined by counting the white colonies on LB+cam+strep+Xgal+20nM 3OC6HSL plates, which distinguish the *S*. Typhimurium signaler from the *E. coli* responder by inducing the latter to switch ON and form blue colonies. The responder CFU/g and the % switched ON were determined on the LB+kan+cam+strep+Xgal plates. Samples collected before the oral gavage resulted in no growth on the selective plates, confirming the absence of the background flora resistant to the antibiotics used to select for the engineered cells. Note that we measured the % switched ON up to Day 6 but tallied the CFU/g until Day 9. In analysis, the % blue colonies of 0.1% or above was considered bona-fide switching ON.

## Acknowledgements

We appreciate Dr. James Slauch for the generous gift of the strain JS481; David Lips for help characterizing various quorum sensing systems in a related work; Dr. Losick, Dr. Desai, Dr. Weiss for providing feedback on the project; and Dr. David Riglar, Dr. Bryan Hsu, and Michael Florea for carefully reading the draft and providing helpful feedback. This work was supported by Defense Advanced Research Projects Agency grant HR0011-15-C-0094.

## Author contributions

PAS, SK, JCW set the general direction of the study; SK designed and conducted all experiments except building and integrating the *S*. Typhimurium memory element (SJK) and making the SPI-2 knockout (MZ); SK analyzed the data with the guidance of PAS; SK built the models; LB, GG, and JCW provided guidance on the animal experiments to study bacterial interactions; SK and PAS prepared the manuscript.

## Declaration of interest

The authors declare no competing interests.

---

**Supp Fig 1.**
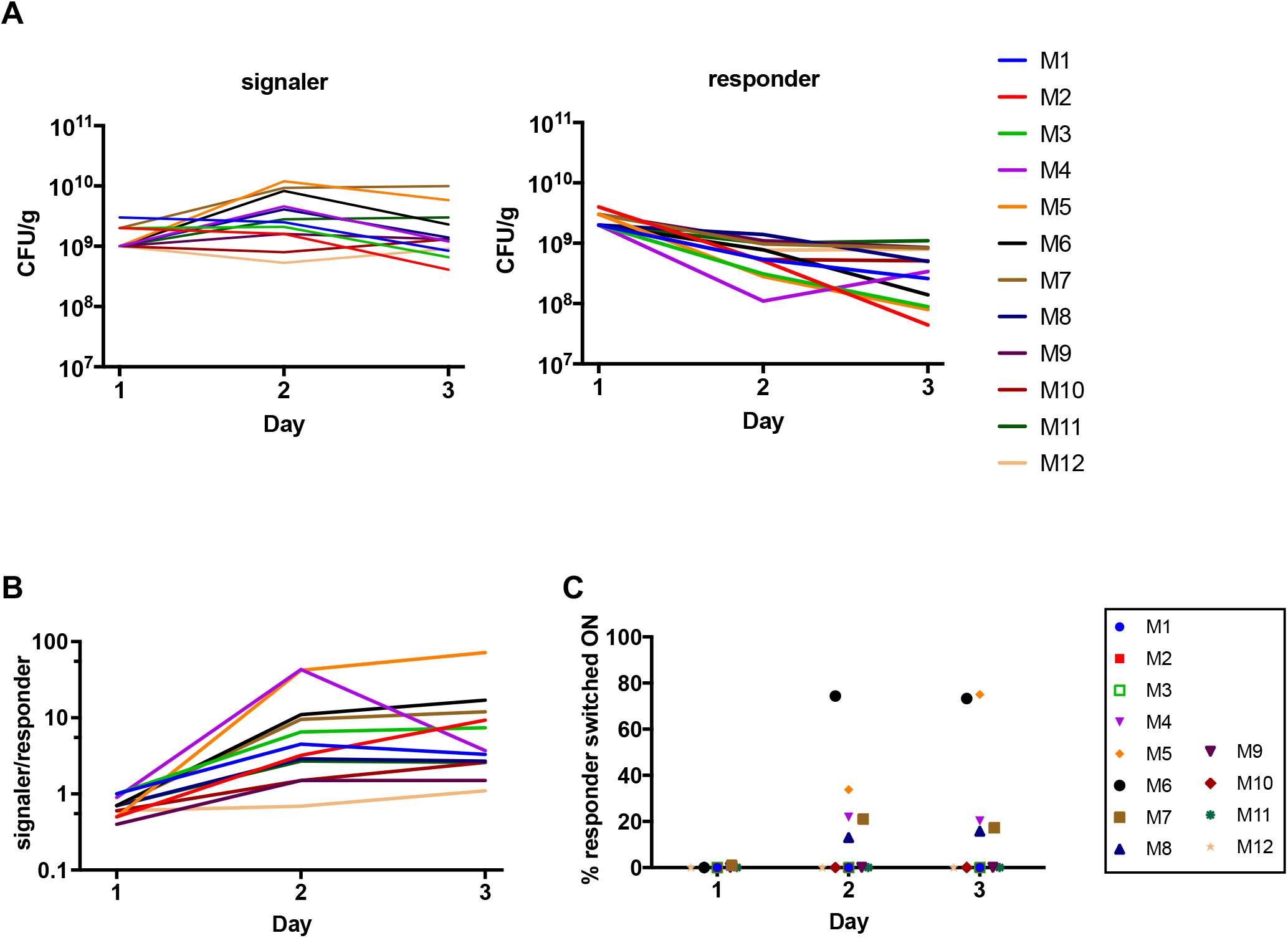
Fluctuations and % switched ON are not correlated in the intra-species information transfer between the *E. coli* signaler and the *E. coli* responder in the mouse gut. (A) Fluctuation in the CFU/g values of the *E. coli* signaler and the *E. coli* responder (B) Fluctuation in the signaler/responder ratios (CFU/g over CFU/g) (C) % responder switched ON of individual mice. M1-M8 are in the ATC group and received ATC water starting after the sample collection on Day 1; M9-M12 did not receive ATC.

**Supp Fig 2.**
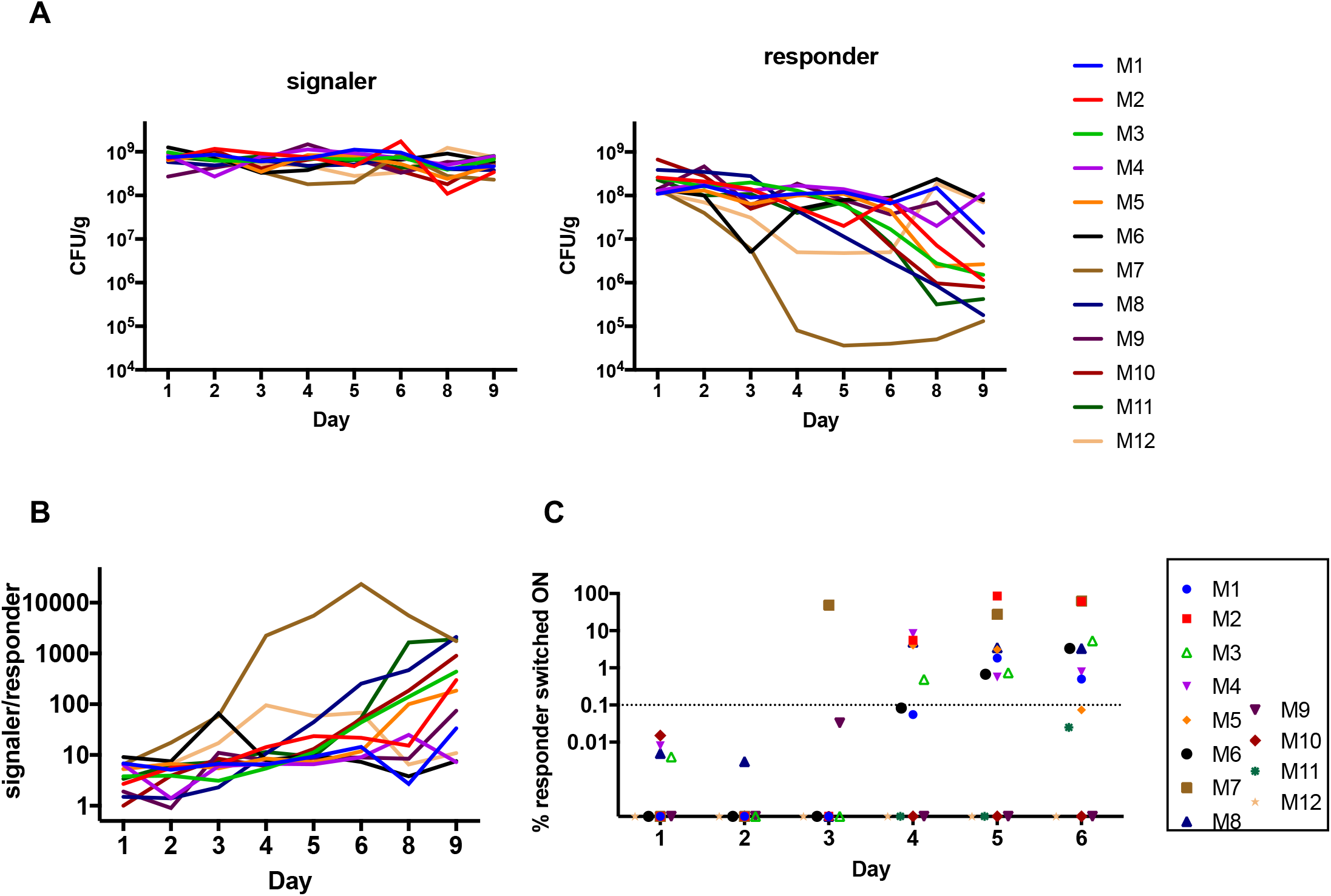
Fluctuations and % switched ON are not correlated in the inter-species information transfer between the *S*. Typhimurium signaler and the *E. coli* responder in the mouse gut. (A) Fluctuation in the CFU/g values of the *S*. Typhimurium signaler and the **E. coli** responder (B) Fluctuation in the signaler/responder ratios (CFU/g over CFU/g) (C) % responder switched ON of individual mice. M1-M8 are in the ATC group and received ATC water starting after the sample collection on Day 3; M9-M12 did not receive ATC.

